# Mutation Maker, An Open Source Oligo Design Platform for Protein Engineering

**DOI:** 10.1101/2020.06.26.171819

**Authors:** Kaori Hiraga, Petr Mejzlik, Matej Marcisin, Nikita Vostrosablin, Anna Gromek, Jakub Arnold, Sebastian Wiewiora, Rastislav Svarba, David Prihoda, Kamila Clarova, Ondrej Klempir, Josef Navratil, Ondrej Tupa, Alejandro Vazquez-Otero, Marcin W. Walas, Lukas Holy, Martin Spale, Jakub Kotowski, David Dzamba, Gergely Temesi, Jay H. Russell, Nicholas M. Marshall, Grant S. Murphy, Danny A. Bitton

## Abstract

Protein engineering is the discipline of developing useful proteins for applications in research, therapeutic and industrial processes by modification of naturally occurring proteins or by invention of *de novo* proteins. Modern protein engineering relies on the ability to rapidly generate and screen diverse libraries of mutant proteins. However, design of mutant libraries is typically hampered by scale and complexity, necessitating development of advanced automation and optimization tools that can improve efficiency and accuracy. At present, automated library design tools are functionally limited or not freely available. To address these issues, we developed Mutation Maker, an open source mutagenic oligo design software for large-scale protein engineering experiments. Mutation Maker is not only specifically tailored to multi-site random and directed mutagenesis protocols, but also pioneers bespoke mutagenic oligo design for *de novo* gene synthesis workflows. Enabled by a novel bundle of orchestrated heuristics, optimization, constraint-satisfaction and backtracking algorithms, Mutation Maker offers a versatile toolbox for gene diversification design at industrial scale. Supported by *in-silico* simulations and compelling experimental validation data, Mutation Maker oligos produce diverse gene libraries at high success rates irrespective of genes or vectors used. Finally, Mutation Maker was created as an extensible platform on the notion that directed evolution techniques will continue to evolve and revolutionize current and future-oriented applications.

## Introduction

The process of natural selection has generated a large set of protein sequences with a broad array of biologically important functions (*e.g.* binding, scaffolding, and catalysis). While the set of extant protein sequences is large (>10^6^), it is only a miniscule fraction of all possible protein sequences (Hecht et al., 2018). Indeed, early efforts in protein design and engineering established that natural protein sequences could be modified to possess properties useful outside biological systems and that entirely *de novo* proteins could be created with structures and functions not observed in nature (Dahiyat and Mayo, 1997; Hecht et al., 1990; Kamtekar et al., 1993; Kuhlman et al., 2003; Moore and Arnold, 1996; Regan and Degrado, 1988). Further advances in directed evolution and protein design combined with decreasing costs of DNA synthesis and sequencing have led to a new wave of biological engineering referred to as Synthetic Biology. Synthetic Biology seeks to redesign naturally occurring biological system (e.g. enhancing metabolic pathways to produce natural products) or to engineer *de novo* biological systems (*e.g.* invention of entirely novel metabolic pathways)(Cravens et al., 2019). Central to the efforts of Synthetic Biology is the ability to rapidly assemble novel genes and introduce mutations as part of a large-scale screen and/or selection. Synthetic Biology efforts to engineer metabolic pathways to produce industrially or pharmaceutically important compounds typically require numerous iterative rounds of gene assembly, mutagenesis, and screening. Given the significant resources required to undertake these efforts, new methods for gene assembly, mutagenesis, and screening are of considerable interest.

Over the past 30 years, numerous techniques and commercial kits have been developed to assemble novel genes and systematically mutate genes at random or specific sites (Mate et al., 2016; Neylon, 2004; Qu et al., 2020; Tee and Wong, 2017). Site Saturation Mutagenesis (SSM) is a widely used technique for construction of a gene library in which a single residue is randomly substituted with all possible amino acids at a single or at multiple positions (Arnold et al., 2003; Miyazaki and Arnold, 1999; Siloto and Weselake, 2012). SSM can be combined with a “site-scanning” strategy (Acevedo-Rocha et al., 2018) to form a technique we refer to as Site-Scanning Saturation Mutagenesis (SSSM, Figure 1A). SSSM can generate a diverse gene library with a theoretical diversity that is appropriately scaled to the throughput of the screening method. Briefly, in SSSM, a set of amino acids in a given protein sequence are subjected to random mutagenesis (at a single site out of defined positions per clone). Target sites might be selected by rational design based on sequence, structural or experimental information, or alternatively through an exhaustive scan of the entire sequence. SSSM is typically performed using mutagenic oligonucleotides (oligos) often referred to as primers, which enable parallel and deep mutational scans that often result in the discovery of beneficial single mutations. These single mutations can then be combined by Multi Site-Directed Mutagenesis (MSDM) (Hogrefe et al., 2002), which generates a combinatorial gene library that permits a comprehensive search for additive mutational effects (Figure 1B). Both methods rely on annealing of mutagenic primers to complementary DNA sequences of a template gene, where mismatches at designated positions facilitate codon substitutions. When multiple amino acid substitutions at the same site are being considered, degenerate primers (oligonucleotides synthesized with multiple nucleotides are a particular position) are often employed to reduce the number of primers to be synthesized and consequently to save time and cost. Once gene libraries are constructed, the genes are transformed into a host (mixed with cell free expression reagents) to enable protein to be expressed and screened for function.

**Figure 1.**
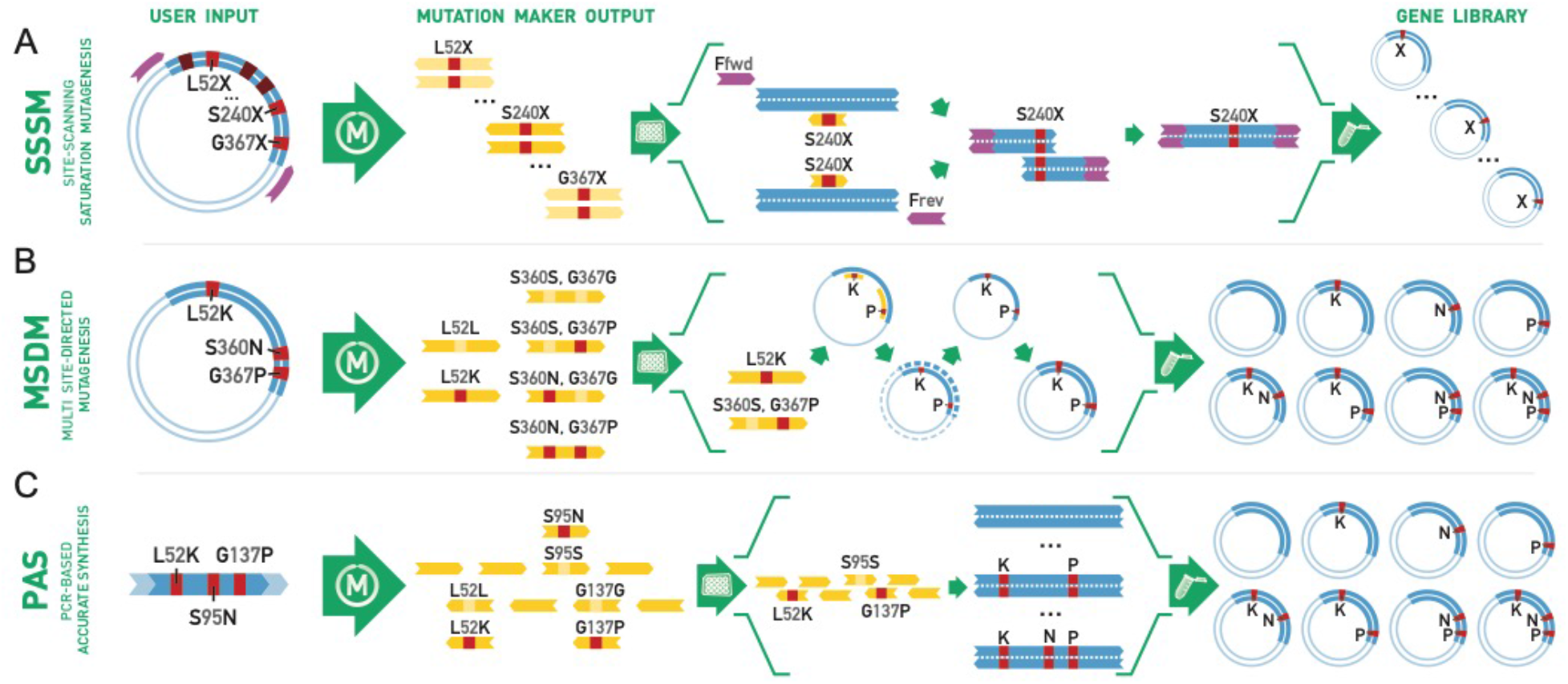
Schematic representations of Site-Scanning Saturation Mutagenesis (SSSM), Multi Site Directed Mutagenesis (MSDM) and Gene synthesis by PCR-based Accurate Synthesis (PAS) workflows. (A left to right) SSSM workflow: user’s input includes plasmid sequence (blue) with gene of interest, a list of amino acid positions to be changed (red squares, only 3 sites are labelled for simplicity), forward and reverse flanking primers (Ffwd and Frev, purple arrows), degenerate codon (marked as ‘X’), and a list of desired parameters (Tm, GC content, flanking primers etc. not shown). Mutation Maker outputs pairs of forward and reverse overlapping mutagenic primers (yellow) for multiple parallel PCR reactions in multiple 96-well plates (green arrow). To create N- and C-gene fragments, PCR reactions are first conducted separately and in parallel for each site (top and bottom rows, respectively) using mutagenic (yellow with red squares) and flanking primer pairs (purple) and a gene template (blue). The overlapping N- and C-fragments are then stitched together using flanking primers (purple) via a second PCR reaction (SOEing, Splicing by Overlap Extension) that produce full-length gene variants. These are subsequently pooled to create a gene library, where each variant carries a single random amino acid substitution at a specified position. Thereafter, the library is cloned into an expression vector (blue circles). (B left to right) MSDM workflow: user input includes gene of interest and a list of target residues and mutations (red squares, only 3 mutations are shown for simplicity) and a list of parameters (Tm, GC, host organism, degeneracy etc. not shown). Mutation Maker outputs a set of unidirectional mutagenic primers (yellow with red squares) as well as primers carrying parental codons (light yellow squares on primers) that cover one or more target sites. The resultant primers can be used in MSDM experiments using the QuikChange Lightning Multi Site-Directed Mutagenesis kit (Agilent) to yield diverse combinatorial library (blue circles). (C, left to right) *De novo* gene synthesis workflow using the PCR-based Accurate Synthesis (PAS) Protocol. The input to mutation maker consists of gene of interest (blue line) with a list of target residues and mutations (red squares, as indicated) as well as and a list of constraints (Tm, GC, host organism, motifs to exclude, degeneracy etc. not shown). Mutation Maker outputs a set of overlapping oligos (yellow) that may carry mutation and parental codon combinations (red and light-yellow squares, respectively) in their non-overlapping portions. These oligos can be assembled to full-length gene variants (blue lines), each carrying different mutation combinations that later can be cloned into an expression vector (blue circles).

In addition to the conventional mutagenesis methods described above, *de novo* gene synthesis methodologies also offer rapid, accurate and cost-effective approaches for assembling and modifying gene sequences (Xiong et al., 2008, 2006). Gene synthesis methods do not require a DNA template and typically rely on synthetic oligonucleotides with overlapping overhangs that can be assembled into a full-length gene by repeated PCR extensions (Figure 1C).

Irrespective of the underlying protocol, the success of mutagenesis and *de novo* gene synthesis experiments is largely dependent on the design quality of mutagenic oligos. To ensure efficient PCR amplification and consequently precise introduction of mutations, a lengthy list of factors must be accounted for during the design phase (e.g. melting temperature (Tm), GC content, primer size, mutation positions, hairpins, primer-dimers, etc.). Several commercial and academic tools were developed to enable design of mutagenic oligos for mutagenesis (Acevedo-Rocha et al., 2018; Camilo et al., 2016; Genee et al., 2015; Novoradovsky et al., 2005) and gene synthesis experiments focused on a limited number of positions or genes. (Bode et al., 2009; Gould et al., 2014; Hoover, 2012, 2002; Richardson et al., 2006; Rouillard et al., 2004; Rydzanicz et al., 2005; Swainston et al., 2014). These tools were not intended to support the scale and complexity of mutagenic oligo design as required by industrial synthetic biology groups. Moreover, none of the available tools offer an automated and customizable design of mutagenic oligos for the construction of combinatorial gene libraries by gene synthesis methodologies. In addition, some of the published tools are no longer being supported, not freely available for widespread commercial use or can handle only a limited number of mutations, if any.

To fill this gap, we developed Mutation Maker, an open source mutagenic oligo design software for mutagenesis and mutagenic *de novo* gene synthesis experiments. Mutation Maker features unique functionalities that are powered by several novel algorithms, providing an accurate, rapid, high-throughput and user-friendly platform for designing large-scale protein engineering experiments. Experimental validation of Mutation Maker revealed high success rates of mutagenic oligos across different genes, vectors and workflows. Mutation Maker workflows were designed primarily to support SSSM using an overlap extension PCR protocol (Horton et al., 1989), MSDM using the QCLM kit (QuikChange Lightning Multi Site-Directed Mutagenesis) (Hogrefe et al., 2002) and gene synthesis using the PAS technique (PCR-based Accurate Synthesis) (Xiong et al., 2006). Oligo design parameters are user-adjustable throughout all gene diversification modalities to allow seamless adoption of alternative and future oligonucleotide-based mutagenesis methods. Additionally, these flexible user inputs render Mutation Maker applicable to both random and rational design strategies, addressing the growing demand for unbiased, focused libraries (Sayous et al., 2020). Finally, Mutation Maker is freely available and open source software, so that the greater scientific community can both utilize the tool without restriction, as well as continue to develop and broaden its scope.

## Results

### SSSM - Mutation Maker offers brute-force or fast-approximation mutagenic oligo designs

In principle, SSSM can reveal the functional consequences of any amino acid change at every position along a protein sequence (Acevedo-Rocha et al., 2018; Qu et al., 2020). SSSM employs multiple parallel PCR reactions that utilize overlapping mutagenic primer pairs and a SOEing PCR technique (Splicing by Overlap Extension) (Horton et al., 1989) to introduce random mutations at specified locations (Figure 1A). To maximize the success of the numerous parallel PCR reactions that are typically performed using multiple 96-well plates, we developed two constraint-satisfaction algorithms, as follows. First, we developed a brute-force algorithm by which all potential primers around all mutation sites are precomputed, and thereafter systematically filtered to satisfy various constraints. Second, we built a fast-approximation algorithm by which mutagenic primers are dynamically extended until they satisfy the design criteria.

The minimal input for both SSSM algorithms is the plasmid sequence containing the parental Gene of Interest (GOI), the forward and reverse flanking primers and a list of mutation sites with their respective degenerate codon (default, “NNK”). Degenerate codons can be directly specified by the user or alternatively can be dynamically generated by a heuristic algorithm from a user-specified lists of amino acids, as described later.

To mitigate known primer-related issues such as primer self-annealing, primer-dimers, and low amplification efficiency, users can adjust the following primer parameters: hairpins and primer-dimers check, GC content, primer length, 5’ end length, 3’ end length, 3’ Tm range, and the overlapping segment size and its Tm range. Since Tm consistency across all parallel reactions (on each 96-well plate) is desired and essential to consistent high-quality PCR, users can also set the maximum Tm difference allowed across the entire set of primers and overlaps.

Both SSSM algorithms consider the lengths of the primer and its 3’ end as hard constraints that must be satisfied, while all remaining parameters are considered soft and may be violated. Mutation Maker also incorporates a scoring function with user-adjustable weights, which governs the logic of constraints violation and consequently of primer ranking. Once designed, the program displays the primers and their respective scores and notifies the user of the violated constraints.

In summary, Mutation Maker factors in multiple user-specified constraints to ensure flexible and reliable mutagenic primer design. It also provides a choice between a fast-approximation search and an exhaustive scan, which in essence reflects a tradeoff between a shorter computation time and a higher probability of finding a better solution, respectively.

### SSSM-Searching for the optimal primer set using a novel brute-force approach

Brute-force algorithms are typically slow, since they exhaustively search for the optimal solution among all possible solutions. Mutation Maker provides users with the option to precompute all potential primers in a specific temperature range and systematically search for the optimal primer set that fulfills the design criteria (Supplementary Figure S1).

First, the brute-force algorithm precomputes all possible forward and reverse primers around each mutation site with or without the use of Primer3 (Rozen and Skaletsky, 2000; Untergasser et al., 2012). The resultant forward and reverse primers are filtered based on their 3’ end length and are paired based on their overlapping region size. Next, the algorithm iteratively loops through the user-specified 3’ end and overlap Tm’s, and for each Tm combination and for each mutation site it selects the best-scoring primer pair. The primer score is computed from deviations between true and expected values of the 3’ end and overlap Tm’s, GC content, and the minimum 3’ end length. Depending on user input, the score function may also penalize primers for hairpins, homo- and hetero-dimers. Lastly, for each set of primer pairs in each temperature step the algorithm computes a cumulative solution score and ultimately reports the best scoring primer set as its best solution.

### SSSM - Novel fast-approximation algorithm generates highly efficient mutagenic primer set

In contrast to the exhaustive approach that precomputes all potential primers for all mutation sites, the fast-approximation algorithm dynamically grows the primers in its search for a mutagenic primer set that least violates the design criteria (Supplementary Figure S2).

The approximation algorithm first iterates through a user-specified range of primers’ overlap Tm’s and for each mutation site it finds the shortest primers’ overlap with a Tm that satisfies an arbitrary threshold. Then, it loops through a temperature grid that is derived from the user’s input and it iteratively grows the forward and reverse primers (one base at a time) from the shortest overlap outwards until the desired 3’ end Tm’s are reached. Similar to the brute-force approach, this algorithm scores and ranks the primers, generating multiple candidate solutions, each containing a complete set of primer pairs for all mutation sites. The algorithm concludes by computing a cumulative score for each solution that ultimately determines the identity of the best solution.

An *in-silico* comparison between the two SSSM algorithms revealed that primer sets produced by the fast-approximation method deviates significantly more from the optimal design criteria compared to the brute-force approach (p < 0.05; Figure 2A). Despite the computationally determined differences, laboratory results clearly indicate that the fast-approximation method generates functional primer sets with high success rates (on average 79.9%) across multiple genes, vectors and host cells (Table 1). Success rates fell within the expected efficiency range of amplification with Q5 Hot Start High-Fidelity DNA polymerase and gene assembly kit according to the product manual (New England Biolabs). Given its significantly shorter computation time (Figure 2B; p <0.05) and its high success rate in laboratory testing, we set the fast-approximation method as the default algorithm for SSSM.

**Figure 2.**
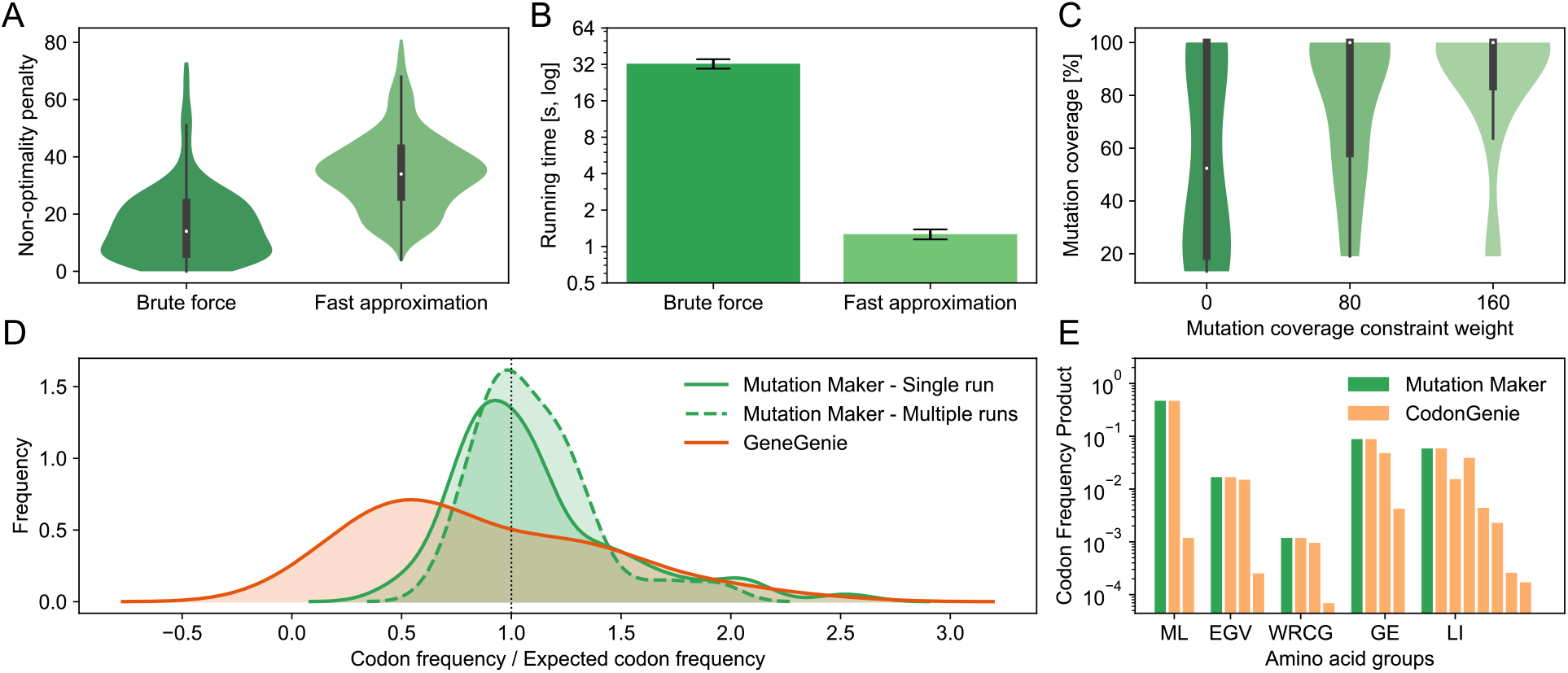
Distributions of (A) Site-Scanning Saturation Mutagenesis (SSSM) cumulative design penalties (log_10_) for brute-force (dark green) and fast-approximation (light green) algorithms (B) SSSM processing times (log_10_ seconds) for brute-force and fast-approximation designs (dark and light green, respectively). 50 Genes with 10 mutations were simulated. Wilcoxon Rank-sum was used to assess the difference between the two distributions (p ~ 6.19e-92). (C) Distribution of Mutation coverage for Multi-Site Directed Mutagenesis (MSDM) workflow as a function of user-adjustable weights. A total of 10 Genes with 2 to 10 mutation sites with up to 5 mutations per site were simulated. X-axis (user-defined mutation coverage weights: 0, 80 and 160 dark to light green); Y-axis (percentage coverage of specified target sites) (D) Distribution of ratios of observed frequency over expected frequency of codons in protein sequences that were reverse translated and optimized by Mutation Maker CFP (Codon Frequency Product) and GeneGenie CAI (Codon Adaptation Index) codon optimization methods. A total of 5 random protein sequences (250 amino acids in length) were reversed translated by Mutation Maker, once (green, solid) or multiple times (green, dashed) and compared to GeneGenie’s (orange). Mutation Maker’s reverse translation incorporates randomization, therefore every protein sequence was reverse translated 5 times and results were summarized as shown. (E) Codon Frequency Product (log_10_ CFP) of degeneracy codons designed by Mutation Maker (green) and CodonGenie (orange) degeneracy algorithms. Random groups of amino acids were used as indicated. Note that CodonGenie outputs multiple solutions whereas Mutation Maker only reports one degenerate solution with the highest CFP.

**Table 1.** Success rates of Site-Scanning Saturation Mutagenesis (SSSM) primer sets designed by Mutation Maker’s novel fast-approximation approach for gene X and gene Y (~700 bp and ~1.2 kbp). Libraries were cloned into different *E. coli* strains using different vectors as indicated. Clones were randomly picked and validated using Sanger Sequencing.

| Library | Number of target sites | Clones tested | Vector | Host Cell | Success Rate (%) |
| --- | --- | --- | --- | --- | --- |
| Gene X -1 | 96 | 48 | pAVE011-Kan | W3110 | 77.1 |
| Gene X -2 | 95 | 24 | pAVE011-Kan | W3110 | 75 |
| Gene X -3 | 31 | 24 | pAVE011-Kan | W3110 | 91.2 |
| Gene X -4 | 96 | 24 | pET30a(+) | BL21(DE3) | 78.3 |
| Gene Y -5 | 96 | 16 | pET30a(+) | BL21(DE3) | 81.3 |
| Gene Y -6 | 96 | 16 | pET30a(+) | BL21(DE3) | 81.3 |
| Gene Y -7 | 96 | 24 | pET30a(+) | BL21(DE3) | 75 |
|  |  |  |  | **Average** | **79.9** |

### MSDM - combining specific amino acid changes using fast-approximation algorithm

Once beneficial mutations are identified by SSSM, scientists often seek to combine these single mutations in order to identify synergistic interactions that may enhance protein performance even further. Widely used commercial kits such as QCLM (Hogrefe et al., 2002) offer simple and rapid protocols to perform MSDM. Briefly, QCLM generates combinatorial gene library via a multi-step protocol that involves template denaturation, mutagenic primer annealing and extension, enzymatic DNA repair and template elimination and transformation (Figure 1B).

The minimal input for the MSDM algorithm is the GOI and a list of target residues alongside their respective amino acid changes. Users may specify the 5’ and 3’ flanking sequences to account for mutations at extreme gene ends. Similar to SSSM, the following primer parameters can be adjusted by the user: primer length and Tm intervals, 5’ and 3’ end length intervals, GC content, hairpins and primer-dimers check and Tm consistency threshold across all primers. Users can also select the non-overlapping option to design primers that can be annealed simultaneously. Without such an option, and under default settings, the algorithm may output partially overlapping primers for neighboring mutation sites that can be used only in successive rounds of MSDM experiments. To account for codon usage and to exclude rare codons users can specify the host organisms and a codon frequency threshold, respectively. Among the above constraints, only the lengths of the primer and its 5’ and 3’ ends serve as hard constraints that are never violated, while the remaining constraints are considered soft. Similar to SSSM, users can set the score function weights to control constraints violation and primer ranking. The MSDM workflow designs degenerate primers by default using a novel optimization algorithm described below. Once designed, unidirectional primers are displayed alongside their properties, including their computed ratios in the final reaction mix.

The MSDM algorithm first identifies all possible partitions of mutation sites, where each partition consists of groups of neighboring target sites that can be covered by a single primer, given the input constraints (Supplementary Figure S3). Then, for each mutation group within each partition and at each primer’s Tm iteration, the algorithm combines selected codons that encode for the desired amino acids and computes all possible minimal primers around these mutation sites. Next, the 3’ ends of these minimal primers are then iteratively extended until they reach the desired Tm at each Tm step. Once grown, the algorithm computes scores for all primers in each mutation group and records the best scoring primer set at each iterated Tm. Upon completion of all iterations, the algorithm computes a cumulative solution score for all recorded solutions and ultimately outputs the best scoring primer set as its optimal solution. The MSDM algorithm may report an incomplete primer set that covers only some of the specified mutation sites, hence its cumulative score also accounts for mutation coverage. User-adjustable weights can be used to increase mutation coverage (Figure 2C). Experimental validation of MSDM primer sets revealed high success rates (72%) with an average of 2.5 mutations per clone (Table 2). According to the QCLM kit manual, success rates were in line with the expected mutagenesis efficiency for 3 simultaneous mutations (>55%). To conclude, Mutation Maker’s MSDM workflow outputs quality primer sets that can produce diverse combinatorial gene libraries using the QCLM protocol.

**Table 2.** Success rate of Multi Site Directed Mutagenesis (MSDM) primer sets designed by Mutation Maker’s novel fast-approximation approach for gene X (~700 bp). Libraries were cloned into *E. coli* strain using vector as indicated. Clones were randomly picked and validated using Sanger Sequencing. Low (L), Medium (M) and High (H) refers to the respective primer to template ratios interrogated (see Materials and Methods). *processed by an earlier version of Mutation Maker (development version), few primers were manually adjusted to account for additional mutations.

| Library | Number of target mutations | Vector | Host Cell | Success Rate (%) | Average number of mutations per clone |
| --- | --- | --- | --- | --- | --- |
| Gene X - 1 L* | 15 | pET30a(+) | BL21(DE3) | 68.8 | 2.6 |
| Gene X - 1 M* | 15 | pET30a(+) | BL21(DE3) | 75.0 | 2.9 |
| Gene X - 1 H* | 15 | pET30a(+) | BL21(DE3) | 81.3 | 3.6 |
| Gene X - 2 L* | 18 | pET30a(+) | BL21(DE3) | 61.5 | 2.5 |
| Gene X - 2 M* | 18 | pET30a(+) | BL21(DE3) | 81.3 | 2.0 |
| Gene X - 2 H* | 18 | pET30a(+) | BL21(DE3) | 64.3 | 2.8 |
| Gene X - 3 L | 14 | pET30a(+) | BL21(DE3) | 46.2 | 3.0 |
| Gene X - 3 M | 14 | pET30a(+) | BL21(DE3) | 71.4 | 1.2 |
| Gene X - 3 H | 14 | pET30a(+) | BL21(DE3) | 43.8 | 2.1 |
| Gene X - 4 M | 14 | pET30a(+) | BL21(DE3) | 87.5 | 2.9 |
| Gene X - 5 L | 13 | pET30a(+) | BL21(DE3) | 100 | 2.1 |
| Gene X - 5 M | 13 | pET30a(+) | BL21(DE3) | 81.3 | 2.5 |
| Gene X - 5 H | 13 | pET30a(+) | BL21(DE3) | 73.7 | 2.6 |
|  |  |  | **Average** | **72** | **2.5** |

### De Novo Gene Synthesis - A Bundle of Novel Algorithms Enables Mutagenic Oligo Design

PCR-based gene synthesis protocols such as PAS rely on short synthetic oligonucleotides that overlap one another by approximately 21 bases and have similar length (~60 bases) and overlap Tm (~56°C) (Xiong et al., 2006). The overlapping oligos cover the entire gene sequence and can be assembled into a full-length gene by high-fidelity PCR reactions (Figure 1C).

Although the principle underpinning gene synthesis is straightforward, computational design of overlapping oligos in practice can be challenging, especially the design of mutagenic oligos for gene diversification purposes. To ensure success of mutagenic gene synthesis experiments, oligos must exhibit unified length and Tm and mutations should not reside within their overlapping segments. In addition, other factors such as the sequence type (protein or DNA), mutation ratios, degeneracy, codon usage, and exclusion of certain motifs also contribute to the complexity of the design task. To address this challenge, we developed a bundle of novel interconnected optimization, constraint-satisfaction and backtracking algorithms that together find mutagenic oligos with unified length and Tm that also satisfy a multitude of user-defined criteria. As elaborated below, the *de novo* gene synthesis workflow incorporates (i) a reverse translation algorithm that optimizes codon frequencies with respect to natural frequencies of the host organism (ii) a constraint-satisfaction algorithm with a backtracking component that splits a given gene sequence into an even number of overlapping DNA fragments that conform to length, Tm, mutation location, GC content and motif exclusion constraints (iii) a recursive degeneracy algorithm that outputs degenerate oligos.

### De Novo Gene Synthesis-Solving Reverse Translation Through Rigorous Codon Optimization

To initiate overlapping oligo design for gene synthesis, users may input the GOI in a DNA or a protein sequence format. In case of the latter, reverse translation must be performed with respect to the native codon frequencies of the host organism to ensure efficient translation. Unfortunately, the level of inherent degeneracy in the genetic code makes reverse translation a non-trivial optimization problem, where a given protein sequence could give rise to numerous DNA sequences. To solve this problem, we developed a simplified optimization algorithm that softly optimizes for higher codon frequencies and considers GC content and motif exclusion. Briefly, Mutation Maker’s reverse translation algorithm generates a DNA sequence from a set of randomly selected codons of the host organism (excluding rare codons). Thereafter, for that sequence the algorithm computes the Codon Frequencies Product (CFP) which is proportional to Codon Adaptation Index (CAI, see Materials and Methods) (Sharp and Li, 1987). Then the algorithm verifies that the sequence meets the design criteria in terms of GC content and motifs exclusion. If so, the algorithm appends the current solution to the list of possible solutions and records the highest CFP. These three basic steps are repeated until CFP cannot be significantly improved. Eventually, the solution with the highest CFP is reported as the optimal reverse translated sequence.

A comparison between Mutation Maker’s CFP optimization algorithm and GeneGenie’s CAI optimization (Swainston et al., 2014) revealed a tighter distribution of codon frequencies around the natural frequencies of the host (Figure 2D). We therefore conclude that Mutation Maker’s CFP optimization algorithm better reflects the natural frequencies in a given host organism.

### De Novo Gene Synthesis-Constraint Satisfaction & Backtracking Algorithms Produce Functional Mutagenic Oligos

The ultimate goal of the *de novo* gene synthesis algorithm is to split a given gene sequence into an even number of overlapping fragments that have consistent length and overlap Tm properties (Supplementary Figure S4). To do so, the algorithm iterates through a user-specified overlap Tm range and first groups the mutations that must reside on the same fragment. Then, around these mutation groups the algorithm generates minimal upstream and downstream sequences that mark the boundaries for gene fragmentation and can potentially serve as overlaps between consecutive gene fragments. The algorithm does so by growing these extensions until they satisfy the current overlap Tm and size constraints. At the next step, (while considering these fragmentation boundaries) a backtracking algorithm optimizes the split of the gene by optimizing the positions and the lengths of gene fragments until the entire gene sequence is covered. Briefly, starting from the 5’ end of the gene the backtracking algorithm adds one gene fragment at the time, placing it at the first possible position given the problem constraints. When the next added fragment cannot be placed due to constraint violation the algorithm shifts the last placed fragment. If the last fragment cannot be shifted anymore (due to constraints violation) the algorithm discards the last fragment and tries to shift the previous fragment and so on. Whenever the algorithm reaches a feasible solution that covers the entire gene sequence, it computes a complete solution score and records the highest. At each Tm iteration, the algorithm continues to search for the optimal gene split until it is found or until a timeout limit is reached. Once the algorithm completes iterating through the entire overlap Tm range, the best scoring set of fragments is selected. Next, the algorithm generates mutagenic oligos from these fragments by introducing and combining mutated or degenerate codons that respects the motifs exclusion constraint. At the very end, the algorithm computes the oligo ratios in the reaction mixture that are necessary to achieve the user-defined mutation frequency in the final library.

Degenerate oligos designed by Mutation Maker were successfully used to construct a full-length gene of 1.2Kb in length. A total of 32 overlapping fragments were designed and tested, of which 6 carried 16 mutations at 8 different sites (Figure 3A). The 32 overlapping oligos were split to 3 parts (segments A1, A2, and A3 consisting of 8, 12, and 12 fragments, respectively), or 4 parts (segments B1, B2, B3, and B4 consisting of 8 fragments each) for the first assembly PCR. The first PCR produced the expected gene segments for 8 fragments assembly (A1, and B1-B4) (300~340 nucleotides in length), but not for 12 fragments assembly (A2 and A3, >460 bases, Figure 3B). The second PCR produced full-length gene for both splits (A1-A3 and B1-B4), with or without gel purification and irrespective of the split strategy employed (Figure 3C). Together, indicating that small amount of successful 1^st^ PCR product was sufficient to serve as a template for the 2^nd^ SOEing PCR, even though it was not detectable on agarose gel. Sequencing analysis revealed that all of the clones (32/32) corresponded to the full-length gene.

**Figure 3.**
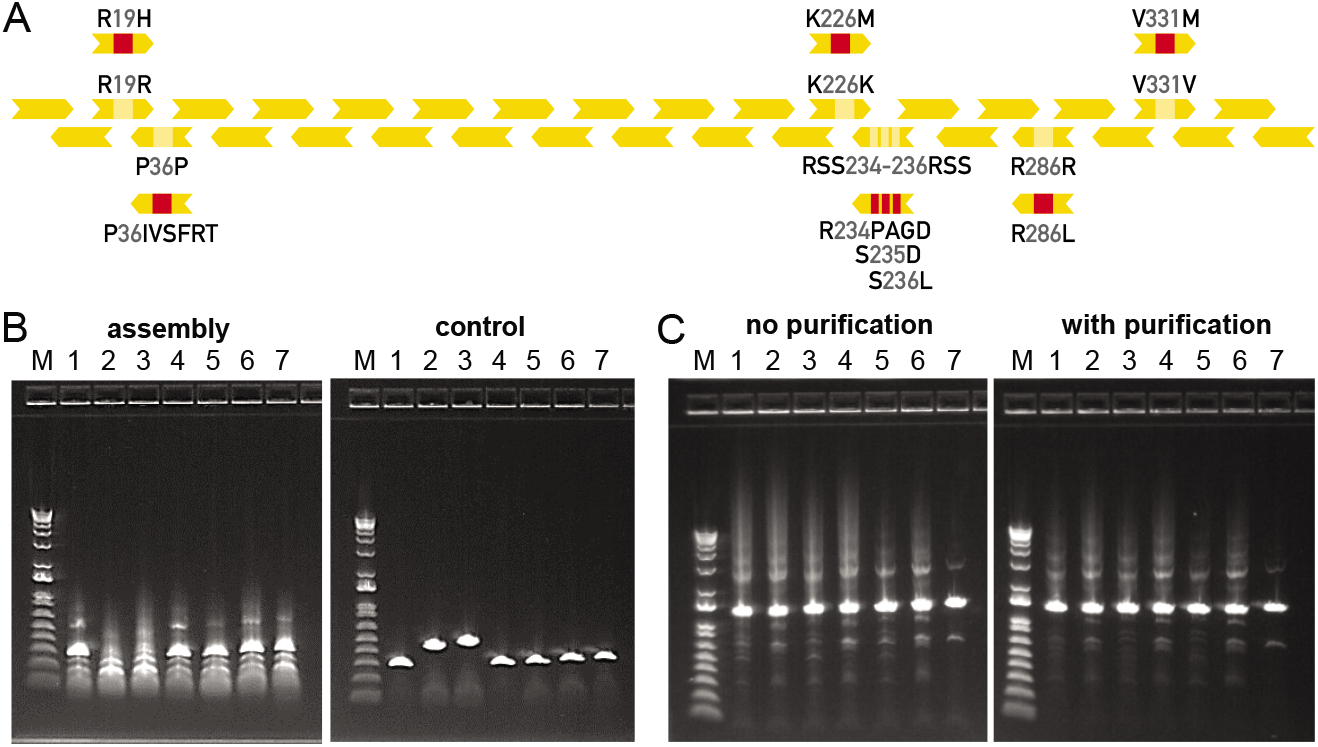
Experimental validation of mutagenic oligos for gene synthesis by PCR-based Accurate Synthesis (PAS) workflow. Degenerate oligos designed by Mutation Maker were successfully used to construct a full-length gene of ~1.2 kbp in length. (A) Schematic representation of 32 overlapping fragments (yellow arrows 5’ and 3’ directions) designed by Mutation Maker, of which 6 fragments carried 16 mutations (red squares) at 8 different sites (as indicated). A total of 50 oligos were used to construct the gene library. (B left to right) The first PCR to create gene segments. Assembly gel: Lane M, 1kb plus DNA ladder (ThermoFisher Scientific); lane 1, segment A1 (8 fragments, 309 bp); lane 2, segment A2 (12 fragments, 466 bp); lane 3, segment A3 (12 fragments, 497 bp); lane 4, segment B1 (8 fragments, 309 bp); lane 5, segment B2 (8 fragments, 315 bp); lane 6, segment B3 (8 fragments, 328 bp); lane 7, segment B4 (8 fragments, 338 bp). Control gel: controls corresponding to the segments form the assembly gel. (C left to right) The 2^nd^ PCR to create full-length gene. Gel without purification of segments from the 1^st^ PCR (no purification): Lane M, 1 kb plus DNA ladder; lane 1, segments A1+A2+A3 (3 PAS); lane 2, segments B1+B2+B3+B4 (4 PAS); lane 3, segments A1+A2c+A3 (2 PAS + 1 control); lane 4, segments B1+B2c+B3+B4 (3 PAS + 1 control); lane 5, segments A1c+A2c+A3c (3 controls); lane 6, segments B1c+B2c+B3c+B4c (4 controls); lane 7, full-length control. Gel with agarose gel purified segments from 1^st^ PCR (with purification) representing the same combination of segments as in the gel with no purification.

Taken together, the gene synthesis workflow features a bundle of novel algorithms for the design of mutated gene fragments that can be assembled into full-length gene variants and generate combinatorial gene libraries.

### Novel Degeneracy Algorithms Solve Different Set Cover Problems

Given the scale of mutagenesis and gene synthesis experiments, scientists often seek to reduce the number of reactions in order to simplify the experimental procedure and lower its overall cost. One way of achieving that is to design degenerate primers that can introduce multiple mutations at a given site via a single reaction. Degenerate primers are essentially a mix of similar oligo sequences that only differ in specific positions and consequently give rise to a population of primers that cover a range of codon combinations (Iserte et al., 2013). Finding an optimal set of degenerate primers corresponds to a set cover problem, where the task is to cover a finite number of elements (amino acids) with as few as possible sets (represented by degenerate codons). Here, we developed two different degeneracy algorithms for two similar yet distinct set cover problems.

In the SSSM workflow, users can directly enter the desired degenerate codons or alternatively can select a set of amino acids that must be included in and/or must be excluded from the degenerate solution. Importantly, if users do choose to compute degenerate codons from inclusion and/or exclusion lists, amino acids that are not explicitly specified may also be included in the degeneracy solution, while only stop codons are excluded by default. To solve this first set cover problem, we developed a heuristic algorithm that first transforms the list of desired amino acids into a set of permuted codons, with one codon per amino acid (maximum number of permutations is predefined). Thereafter, the algorithm iterates over the list of permuted codons, while attempting to identify degenerate alphabets that encode the amino acids specified in the inclusion but not in the exclusion list. If no solution is found, the algorithm splits the list of codons into two subsets and repeatedly searches for a minimal set of degenerate codons in each subset. It does so recursively until the solution satisfies the user input. More specifically, for every iteration, the algorithm updates the current best solution, if the number of codons that are encoded by the degenerate alphabet is lower, and if the degenerate alphabet itself is shorter. The algorithm may report the results before it loops through all possible permutations, when the predefined time limit is reached or when it finds the optimal solution (e.g. finds one degenerate codon that encodes exactly one codon per one amino acid from list of desired amino acids).

In MSDM and gene synthesis workflows the designed degeneracy solution is strictly restricted to the list of specified substitutions in the input. In addition, the solution must encode the parental amino acids at each site and may exclude rarely used codons. To solve this second set cover problem, we developed another heuristic algorithm that randomly selects codons for a given set of amino acids, computes a single degenerate codon for this set and decodes it back to the list of amino acids. This step is repeated multiple times and results are recorded. Then the algorithm retains only degenerate solutions that produced the lowest number of amino acids. From the resultant list, it then selects the solution with the highest CFP. The algorithm then checks whether the generated amino acids list matches the user input. If so, solution is reported. If not, the algorithm splits the set of amino acids into all possible subsets, and recursively repeats the previous steps for each subset, until the degeneracy codon fulfils the design criteria.

Importantly, heuristic algorithms are optimised for computational performance and by definition find a solution that might not be the optimal one. Hence, we next evaluated the performance of our degeneracy algorithm in terms of producing minimal sets of degenerate codons with high CFP. In this respect, Mutation Maker’s degeneracy results were consistent with the highest scoring CodonGenie (Swainston et al., 2017) degeneracy output (Figure 2E). Together we conclude that Mutation Maker enables design of degenerate primers that simplify the experimental procedure and consequently save time and cost.

## Discussion

Protein Engineering will continue to transform the research landscape of biotherapeutics and industrial chemical processes. The ability to generate novel biotherapeutics with designer properties and to develop novel (Schmid et al., 2001; Sheldon and Woodley, 2018) enzymatic processes can be primarily attributed to decades of advancements in protein design and directed evolution techniques that provide new means to optimize protein performance or to create entirely new proteins (Arnold, 2018, 2015; Turner, 2009) (Chari & Hecht). Protein Engineering techniques typically begin with a diverse library of gene variants in a test tube, followed by expression in a host organism and screening or selection for variants with desired properties. Gene diversification is therefore a key step that determines the quality of the library and consequently the success of identifying variants with beneficial mutations at a reasonable effort, time and cost. However, the scale and the complexity of gene diversification in practice demands the development of computational tools for automated library design that can significantly reduce experimental time and mitigate biases and errors. To address this issue, we developed Mutation Maker, a mutagenic oligo design software for SSSM, MSDM and *de novo* gene synthesis workflows that are among the most frequently used gene diversification methodologies. Such protein engineering workflows are well established in the industry and are considered the most efficient procedures for rapid data generation.

The motivation to create Mutation Maker stemmed from the lack of any protein engineering - oriented application that enables bulk design of mutagenic oligos at industrial scale and supports multiple gene diversification modalities for generating random, directed and combinatorial gene libraries. Although several tools for mutagenic oligo design exist (Acevedo-Rocha et al., 2018; Camilo et al., 2016; Genee et al., 2015; Novoradovsky et al., 2005), they are largely limited, obsolete or not freely available for widespread commercial use. For example, AMUSER (Genee et al., 2015) and HTP-oligo designer (Camilo et al., 2016) were intended for site-directed mutagenesis, yet these tools permit only a single mutation per sequence, do not account for degeneracy and offer limited functionalities. Deepscan (Acevedo-Rocha et al., 2018), stands out from the freely available automated design tools, enabling mutagenic primer design at scale. Deepscan certainly represents a significant step forward for designing random gene diversification experiments. Nonetheless, it exhaustively generates primers for every codon in a given sequence and it still lacks many of the functionalities that Mutation Maker has to offer.

With respect to design of gene synthesis experiments, several tools (Gould et al., 2014) including DNAWorks (Hoover, 2012, 2002) and GeneGenie (Swainston et al., 2014) can effectively split genes into synthetic oligos. However, unlike Mutation Maker these tools were not designed to support large-scale protein engineering experiments. More specifically, DNAWorks is limited in terms of introducing multiple mutations, it requires multiple runs and many manual steps, deeming it unsuitable for high-throughput gene library design tasks. Similarly, GeneGenie (Swainston et al., 2014) does not permit introduction of specific mutations, it only accepts protein sequences and it cannot handle large input sequences. Mutation Maker is therefore, to our best knowledge, the only software that enables automated design of combinatorial gene libraries by gene synthesis methodologies.

To ensure efficient translation of synthetic genes in the host organism Mutation Maker also features reverse translation functionality that is tightly coupled to codon optimization. In this regard several proprietary and freely available codon optimization algorithms have been developed (Chung and Lee, 2012; Gould et al., 2014; Liu et al., 2014; Raab et al., 2010; Swainston et al., 2014; Tuan-Anh et al., 2017). However, many of the published methods optimize the codons in the target gene with respect to the CAI (Sharp and Li, 1987). The CAI reflects the deviation of codon frequencies in a given sequence from those observed in a reference set of highly expressed genes in the target host organism. CAI highly correlates with gene expression, hence it has become a preferred optimization method (Gould et al., 2014). Nevertheless, concern has been voiced that CAI maximization may lead to tRNA depletion and consequently to an increase in translational errors (Chung and Lee, 2012). To mitigate such possibility, Mutation Makers offers an alternative CFP optimization method that is proportional to CAI, but only softly optimizes for higher codon frequency and therefore better reflects the native codons landscape in a given host.

Despite its many unique features Mutation Maker inevitably shares some functionalities with existing tools. However, while certain gene diversification features may be distributed across multiple tools and websites, Mutation Maker provides a ‘one-stop shop’ platform for high-throughput protein engineering experiments. Furthermore, Mutation Maker’s backbone is scalable, container-based and open source, providing a robust foundation for heavy workload, future development and further scope expansion by the greater scientific community.

The inherent complexity and plasticity of the novel algorithms developed here necessitated the development of a simplified user interface that maximizes users’ ability to fine-tune key parameters and constraints in order to adapt to different experimental scenarios and protocols. Streamlining and simplifying the entire library design process was of paramount importance, therefore we conducted an extensive user experience research with protein engineering scientists in order to guide Mutation Maker’s architecture and design. To this end, Mutation Maker’s offers an up to date, dynamic and intuitive user interface that features tooltips, accepts multiple input formats, and includes interactive visualization, printing, export and results sharing functionalities.

The novel algorithms described here were optimized and validated by rigorous *in-silico* simulations, which in turn helped to define the default settings of this tool. In addition, mutagenic oligos for all design workflows were experimentally validated in real-world industrial protein engineering projects. Nonetheless, it is important to note the limitations of these algorithms. For performance optimization reasons, the SSSM brute-force approach is only exhaustive within fairly large (HOW LARGE?) Tm windows that slide along the user-specified Tm range. Thus, in essence, not all possible primers across the specified Tm range are interrogated. Mutation Maker by design does not optimize for the presence of GC clamps throughout, since this constraint often eliminated better scoring solutions. The scoring functions for evaluating primers and solutions have been optimized based on past experience and initial laboratory testing but have not been optimized using exhaustive laboratory testing and therefore are subject to improvement over time. Mutation Maker’s gene split algorithm can be also further optimized, since only simplified scores were used to evaluate fragments during backtracking as a tradeoff for reduced computational time.

The gene synthesis algorithm may not guarantee an optimal solution for complex cases when it hits the hardcoded timeout for each Tm iteration. Another limitation worth noting is that the algorithms for reverse translation and for degeneracy are solving these problems using heuristics and randomization and therefore may not output the optimal solution.

Despite the above limitations, laboratory test results were highly successful. Mutation Maker mutagenic oligos were successfully used to generate diverse random and combinatorial gene libraries. Finally, Mutation Maker is a freely available, open source software that can be deployed as web application on a server or locally. Mutation Maker was designed to be modular and extensible, aiming to encourage scientists to add new methods and protocols that can accelerate protein engineering and synthetic biology research and applications.

## Materials and Methods

### Mutation Maker features scalable, open source and container-based architecture

Mutation Maker is composed of five distinct components as follows: (i) A JavaScript (JS) User Interface (UI) where users can submit oligo design tasks and visualize the results using a customized version of the neXtProt viewer (Paladin et al., 2020; Zahn-Zabal et al., 2019), (ii) A Worker that is assigned by the open source Celery asynchronous task queue, where computation takes place, (iii) An Application Programming Interfaces (API) that provide communication between the UI and the Worker, (iv) An open source Redis in-memory database that is used to temporarily store information about the design task and its respective results, (v) The AWS (Amazon Web Services) Lambda service is also optionally employed to parallelize part of the SSSM computation that utilizes Primer3 binary (Rozen and Skaletsky, 2000; Untergasser et al., 2012). However, Mutation Maker can run on any computer that provides 256MB RAM for the UI, 512MB RAM for the API, and 8GB RAM for the Worker. All the aforementioned services can be executed in a local Python and JavaScript environment or alternatively via a Docker Compose script that executes each service as a Docker container. Note that parallelization of Primer3 under local deployment is possible without the need for an AWS account, yet oligo design run time with Primer3 is rather slow in this case.

### User Interface

The web application is based on React, TypeScript and Ant Design JavaScript libraries. Landing page provides links to the three workflows (SSSM, MSDM and gene synthesis). Each workflow consists of an input form with advanced on-the-go validation and a dynamic result page. Results are displayed using interactive table, modified neXtProt Feature Viewer (Paladin et al., 2020; Zahn-Zabal et al., 2019) and Excel export. Results with input parameters can be exported to PDF or as a temporary sharable link. Designed oligos can be exported as annotated XLSX file. The neXtProt Feature Viewer is JavaScript Library developed by Swiss Institute of Bioinformatics (SIB) to visualize protein sequence features using D3 design (Paladin et al., 2020; Zahn-Zabal et al., 2019). These libraries have been modified to meet the visualization requirements for our three design workflows. The following features were added: GC content graph, primer directionality, mutation alignment and integration with interactive results table and with React user interface.

### Melting Temperature calculations Primer dimer and hairpin checks

Melting temperature was computed using Primer3 Python library (Untergasser et al., 2012) (available at https://pypi.org/project/primer3-py/), with the nearest neighbors Santa Lucia approach (SantaLucia, 1998) and the following settings: calculation_method = “santalucia”, salt_correction = “ owczarzy”, na = 50 (Na^+^ ion concentrations, mM), k = 50 (K^+^ ion concentration, mM), tris = 20 (Tris-HCL concentration, mM), mg = 2 (Mg^2+^ ion concentration, mM), dntp = 0.2 (dNTP concentration, mM), dnac1 = 500 (primer concentration, for PAS was set to simulated concentration of 583 mM), precision = 1 (precision of the temperature calculation in digits after the decimal point). For PAS oligo Tm’s a Tm correction factor of 3 °C was applied.

Primer hairpin, homo- and hetero-dimer temperatures were computed using the same Primer3 Python library (Untergasser et al., 2012). Hairpin and primer-dimer penalties were applied to candidate oligos only when primer-dimer or hairpin formation temperatures were very close to the specified optimal reaction temperature. The hetero-dimer penalty is considered only for SSSM and MSDM workflows, while for PAS hetero-dimer is omitted, given the desired overlap between consecutive oligos. No other mispriming checks were conducted for PAS. In SSSM flanking primers were compared with each candidate mutagenic forward or reverse primer. In MSDM all possible heterodimers are considered, since all unidirectional primers are assumed to be used in a single reaction. For the SSSM brute-force approach option, primer-dimer and hairpin checks are performed by default using Primer3 binary. Note that in order to enable longer primer design for SSSM using Primer3, the maximum primer length in Primer3 was increased from 36 to 60 nucleotides in the libremer3.c library.

### Score functions

All workflows utilize score functions that are based on deviations of primers properties from the optimal values. For SSSM and MSDM the solution score consists of sum of scores for each individual primer and for gene synthesis the solution score consists of averaged score of all fragments and overlaps. The score for a single primer (or a fragment) is computed as a root sum of weighted squared errors for all parameters and can be generalized by this formula:

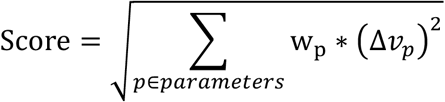

Where:

- p represents iterator over all parameters (melting temperature, primer length, GC content, etc.).
- w_p_ is a user specified weight for the parameter.
- Δv_p_ is the difference of actual parameter value from optimal value.

However, each workflow requires specific amendments to accurately evaluate candidate solution. Detailed description of scoring functions for each workflow are given below.

SSSM Primer Pair score:

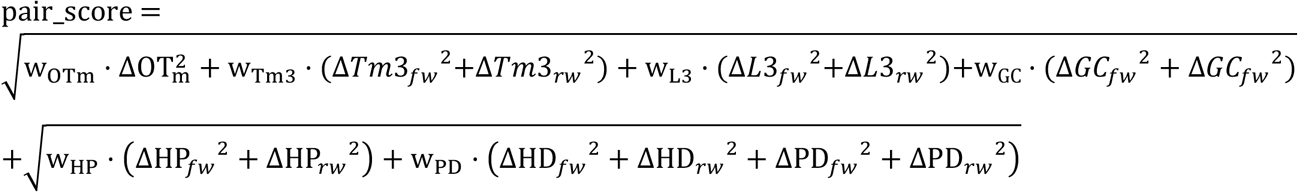

Where:

- ΔOT_m_ is the difference between the melting temperature of overlap and the PCR reaction temperature.
- ΔTm3 is the difference between the temperature of the primer 3’ end and the specified minimal Tm of the 3’ end, for the forward (fw) and the reverse (rw) primers.
- ΔL3 is the difference between the primer 3’ end size and the minimal 3’ end size.
- ΔGC is the deviation of the actual GC content from the specified GC content interval, in percentage.
- ΔHP is the difference between the hairpin melting temperature T_hairpin_ (if the primer can create a hairpin), and the “safe self-binding temperature” T_safe_ which is defined as (PCR reaction temperature) – 2(T_m_ range size). If T_hairpin_ < T_safe_, then this term is ignored.
- ΔHD is calculated the same as ΔHP, but for homodimers.
- ΔPD is calculated the same as ΔHP, but for dimers between the forward primer and the reverse flanking sequence (fw) and between the reverse primer and the forward flanking sequence (rw).
- *wx* are user-supplied weights for each variable *X*(*X* = OT_m_, Tm3, etc.).

SSSM solution score:

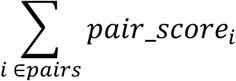

MSDM Primer score:

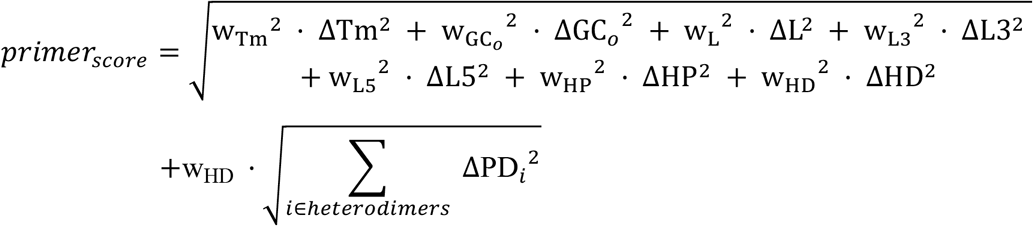

MSDM Solution score:

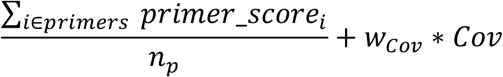

Where parameters are as described for SSSM, with the following differences:

- ΔTm is the difference between the primer melting temperature and the PCR reaction temperature.
- ΔGC_o_ is the difference between the actual and optimal GC content, in percentage.
- ΔL is the difference between the primer length and minimal primer length.
- ΔL5 is the difference between the primer 5’ end size and the minimal 5’ end size.
- ΔPD_i_ is the difference between the melting temperature of the i-th heterodimer (between the current primer and another primer), and the “safe” melting temperature, as in ΔHP used for SSSM.
- *n_p_* is the number of primers.
- Cov is the coverage of mutations by the MSDM solution primers, in percentage.
- Other symbols have the same meaning as in the SSSM scores.

Gene Synthesis Fragment score:

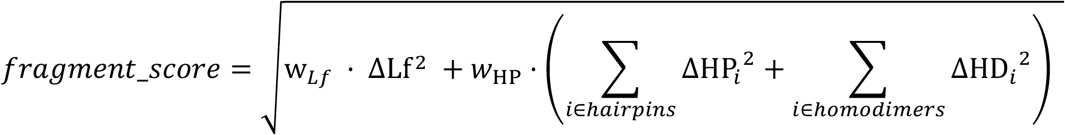

Gene Synthesis Overlap score:

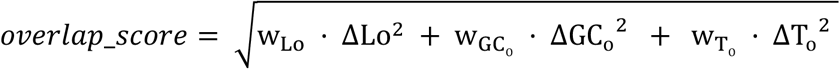

Gene Synthesis Solution score:

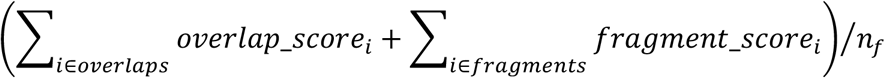

Where:

- ΔL_f_ is the difference between the actual length of a fragment and the optimal length.
- ΔHP_i_ is the value of ΔHP (as described for the SSSM formulas) for the i-th hairpin region in a given PAS fragment. The PAS workflow considers multiple hairpins for the same DNA fragment, as PAS fragments are typically much longer that SSSM or MSDM primers.
- ΔHD_i_ is the value of ΔHD (as described for the SSSM formulas) for the i-th region in a given PAS fragment which can take a part in forming a homodimer.
- ΔL_O_ is a difference between the length of the overlap of two fragments and the optimal length of the overlap.
- ΔGC_O_ is the difference between GC content for the overlap and the specified optimal GC content.
- ΔT_O_ is the difference between overlap temperature and the optimal overlap temperature.
- *n_f_* is the number of fragments.
- Other symbols have the same meaning as in the SSSM scores.

The solution score is the average scores for all fragments and overlaps. The weights used in the score functions are hardcoded. The Mutation Maker uses the following values:

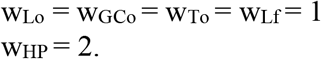

### Backtracking algorithm

The backtracking algorithm of the gene synthesis workflow can be described by the following pseudocode:

~~~
procedure **bt**(c)
  if **reject**(P,c) then return
      else **record**(c)
  if **accept**(P,c) then
      **save_best_solution**(c)
      return
  s ← **first**(P,c)
  while s ≠ NULL do
      **bt**(s)
    s ← **next**(P,s)
~~~

Where:

Input: ‘**c**’ is an ordered list of starts of fragments in a partial solution consisting of DNA fragments covering an initial part of the gene. **P:** is the input data for the gene synthesis algorithm (GOI, constraints etc.) plus selected Tmin for the gene synthesis workflow. **reject**(P, c) is a function that returns true, if new partial solution ‘c’ violates any of the following conditions:

1. Constraints given by P
2. The last fragment from the sequence of ‘c’ must not form a hairpin or homodimer for given Tm.
3. The algorithm has not already found a partial solution that ends at the same nucleotide, has the same parity of the number of fragments (odd, even), and has a lower value of the score function than solution ‘c’.

Condition 3 refers to a dynamic programming optimization, where the score function serves as the optimization criterium and the state is defined by the end position of fragment sequence and parity of the fragment sequence length. Function **record** (c) saves partial solution ‘c’ (list of fragments) and its score in a table with index given by pair (last nucleotide position, parity). The **accept**(P,c) function checks that: (i) the list of fragments ‘c’ covers the whole gene, (ii) the number of fragments is even, (iii) the score of solution ‘c’ is better than the best recorded score for complete solution (constraints ‘P’ are used for computing the score in **accept**()), (iv) the last fragment from ‘c’ does not form a hairpin or homodimer for given Tm. The **save_best_solution**(c) function records sequence ‘c’ and its respective score. Function **first**(P, c) extends partial solution ‘c’ by adding a DNA fragment with the length closest to the optimal fragment length, given constraints ‘P’. Finally the **next**(P, s) function replaces the last fragment in ‘s’ by the next best fragment with respect to the distance from the optimal fragment length that fulfils constraints ‘P’.

### Codon Usage & Product of Codon Frequencies Optimization

To account for species-specific codon usage in the MSDM and gene synthesis workflows Mutation Maker utilizes taxa-specific genetic code tables and species-specific raw codon frequencies. The 33 taxa-specific genetic code tables are available from https://www.ncbi.nlm.nih.gov/Taxonomy/Utils/wprintgc.cgi, while 35,799 species-specific raw sum of codons are available from ftp://ftp.ebi.ac.uk/pub/databases/cutg/ (Nakamura et al., 2000). Codon usage bias is defined as the difference in the frequency of synonymous codons in individual species. These relative codon frequencies can be directly computed from the raw sum of codons in a given species. Mutation Maker’s codon usage function simply uses the standard species taxonomy ID to retrieve the corresponding genetic code table and the respective raw codon frequencies to calculate these relative frequencies.

Mutation Maker’s codon optimization approach softly optimizes for the Codon Frequencies Product (CFP) which is proportional to CAI:

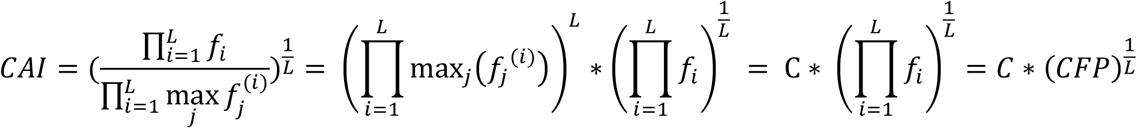

Where 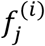 are frequencies for synonymous codons for amino acid i, ‘C’ is a constant for a given gene sequence, and 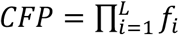.

Better reflected when log is taken for both sides of equation:

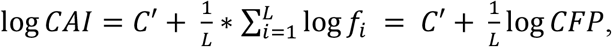

Where *C*’ = log *C*.

### Ratio calculation for gene synthesis

For gene synthesis, users can specify the percentages of occurrence of each mutation in the resulting mixture. This is achieved by setting concentrations of all variants of the DNA fragments generated by the gene splitting algorithm. The desired concentrations of the variants (referred to as “oligos” here) are computed by the following algorithm:

Let *r*_1_^(*i*)^,…, *r*_K_^(*i*)^ be the desired ratios for mutations M_1_,…, M*_K_* at the *i*-th mutation site on the fragment.

We can suppose that:

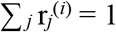

While one can rephrase the problem as follows:

Find a concentration for each oligo, as a [0, 1] share of the total number of molecules, which gives a mixture with the following property: If a molecule is randomly picked from the mixture, the probability that it has mutation M_j_ at the i-th mutation site is equal to r_j_^(i)^. The computation of the oligo concentrations is done according to the following theoretical result:

Suppose that an oligo has mutations M_j1_^(1)^,…, M_jn_^(*n*)^ at mutation sites 1,…, *n*, where *n* is the number of mutation sites on the fragment. Then this oligo share must be equal to *r_j1_*^(1)^..... *r_jn_^(n)^*

Proof:

Let A_1_,…, A_*n*_ be random variables the values of which are mutations at sites 1,…,*n*.

Let’s assume that these variables are *independent*. Then,

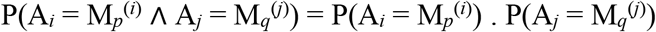

Assuming a random process in which the next oligo to be put in the mixture is repetitively selected from a set of all possible mutated oligos.

If an oligo with a specific mutation combination with probability equal to the product of probabilities for its mutations is selected, then, indeed, the same probability for randomly picking this oligo from the resulting mixture is obtained.

Note that the above statement does not prove that this is the only possibility to achieve the desired mutation ratios. However, it guarantees that this requirement will be satisfied, as long as all possible mutation combinations can be used. For degenerate solution, when an oligo specification can cover multiple mutations at the same site, the solution may not work. In this case, Mutation Maker outputs the concentrations of the affected oligos as an “unknown” value. However, if the ratio of the mutations generated by one degenerate codon is as required (for instance, 1:1), then the same method can be used also for oligos with degenerate codons. If this is the case, then the algorithm just uses a sum of expected ratios for mutations covered by the degenerate codon instead of values for individual mutations as shown in the product above. Another condition that must be met is that the degenerate codons should not overlap for the same mutation site (for instance, degenerate codons for FL and LI should not reside on different oligos for the same site). Otherwise, independence of the random variables as shown in the proof above cannot be assumed.

### Experimental Validation for SSSM

The SSSM primer sets designed by the fast-approximation for gene X and gene Y (~700 bp and ~1,2 kbp in length) were validated as follows. The SSSM library was created by a modified overlap extension PCR method (Horton et al., 1989; Williams et al., 2014). Typically, up to 96 amino acids in GOI sequence were selected as a set of target sites for one library, and forward (‘NNK’) and reverse (‘MNN’) mutagenic primers were designed using the SSSM workflow with default setting. The resultant forward and reverse primers were synthesized and formatted into two 96-well plates separately (Integrated DNA Technology, USA). The GOIs was cloned into pET30a(+) or pAVE011-Kan vector and used as the template for PCR. To create N- and C-terminus mutagenic fragments, the 1^st^ PCR reactions were performed with Q5-Hot Start DNA polymerase (New England Biolabs, USA) using reverse and forward primer plates paired with the 5’ - and 3’ flanking primers, respectively. The following PCR conditions were used: initial denaturation at 98° C for 1 minute; 25 cycles of [98° C for 10 seconds; 60° C for 20 seconds; 72° C for 20-30 seconds/kb]; and final extension at 72° C for 2 minutes. Then, the N- and C-terminus fragments were mixed together in a single 96-well PCR plate and treated with ExoSAP-IT (ThermoFisher, USA) and DpnI to digest the parental plasmid and primers. To create full-length mutant gene, the 2^nd^ PCR (SOEing PCR) reactions were performed using ExoSAP-IT/DpnI treated fragment mixture and 5’- and 3’-flanking primers under the same conditions used for the 1^st^ PCR. After SOEing, amplifications in each well were confirmed by running on 96-well E-gel system (Life Technologies, USA). The mutant GOI fragments were pooled to give a single gene library and cloned back into the plasmid by NEBuilder HiFi assembly kit (New England Biolabs, USA) and transformed to *E. coli* cells. Clones were randomly picked, and the mutations were validated using Sanger Sequencing (GeneWiz, USA).

### Experimental Validation for MSDM

MSDM primer sets designed by Mutation Maker were validated as follows. Gene X (~700 bp in length) was cloned into pET30a vector. The library targeted combinations of approximately 13 to 18 mutations ranging between 6 and 12 sites per library. The MSDM library was created using the QuikChange Lightning multiple-site-directed mutagenesis kit (Agilent Technologies, USA). In short, site-directed primers were designed using MSDM workflow with default settings. The resultant primers were synthesized by Integrated DNA Technology (USA). The primers encoding mutations were mixed in a single tube and used for QuikChange reaction. To confirm the robustness of the protocol, the reaction was performed under three different conditions; Low (L), Medium (M) and High (H), in which 6, 15, and 30 nmol of mutagenic primer mix were used, respectively, against 30 ng of the template plasmid in 15 uL reaction volume. The resultant libraries transformed into *E. coli* BL21(DE3) cells, and isolated colonies were tested by Sanger sequencing (GeneWiz, USA) to confirm library quality.

### Experimental Validation Gene Synthesis (PAS)

The complete synthetic gene library of the test GOI (gene Y, ~1.2 kbp) was constructed using the PAS protocol described by Xiong *et.al.* (Xiong et al., 2006). Briefly, Mutation Maker PAS workflow (under default settings) split the ~1.2 kb gene into 32 overlapping fragments, of which 6 fragments carried 16 mutations at 8 target sites, producing 50 oligo primers in total (Figure 3C). The 1^st^ PCR assembly of the 8 or 12 fragments and the 2^nd^ SOEing PCR assembly of 3 or 4 sub-fragments were performed using Q5-Hot Start DNA polymerase (New England Biolabs, USA). Following each PCR round, the expected bands were purified by agarose gel extraction using ADB buffer and DCC-5 column (Zymo Research, USA). The final library fragments were cloned into pET30a(+) vector using NEBuilder HiFi Gene Assembly kit and transformed *E. coli* BL21(DE3) cells, then the isolated colonies were tested by Sanger sequencing (GeneWiz, USA) to confirm library quality.

### Data Availability

All code for this publication is available in the following GitHub repository: https://github.com/Merck/Mutation_Maker.

## Acknowledgments

We thank Keith A. Canada, Sandra K. Tremps, Carol Rohl, Farida Kopti, Martin Ryzl, and Jens Christensen for their continuous support of this project. We thank Xinnian Li for his help in gathering the project requirements. We are also immensely grateful to Jiri Marsicin and Jan Tkacik who initiated development of the tool and to Jan Barta for his help implementing some of the user interface features. We also thank Christina Ferraro and Wai Ling Cheung-Lee for their help with the experimental validation. We would like to acknowledge and thank Jyoti Shah, Geoffrey Hannigan and Victor Neduva for reviewing an earlier version of the manuscript.

## Author Contributions

DB and KH conceived the study. DB supervised the study. AG gathered the requirements from the scientists and coordinated the project. PM led the development and designed the MSDM and gene synthesis algorithms and JA designed the SSSM workflows. PM, JA, NV and MM implemented the algorithm and wrote the associated tests. AV wrote the gene synthesis simulation tests. NV designed and implemented the reverse translation and degeneracy algorithms for MSDM and gene synthesis and implemented oligo generation and ratio calculation for PAS. RS designed and implemented the degeneracy algorithms for SSSM. LH conducted the user experience research and designed the user interface. SW, MW, RS, MM, DP and JA implemented the user interface. MM, JA, OK, MS, and DP packaged the software and MS prepared it for open source. PM, MM, JN and OT designed and implemented the score functions throughout. KC implemented the multi-species codon frequencies functions. JK, GT, DD, NM, and JR advised throughout the study. KH, DP, DD, MM, OK and DB designed the figures. DP did the artwork. DB wrote the manuscript. PM, MM, NV, KH, KC, SW and GM also contributed to the final draft, all authors read and approved the final version of the manuscript.

## Supplementary Material

**Supplementary Figure S1.**
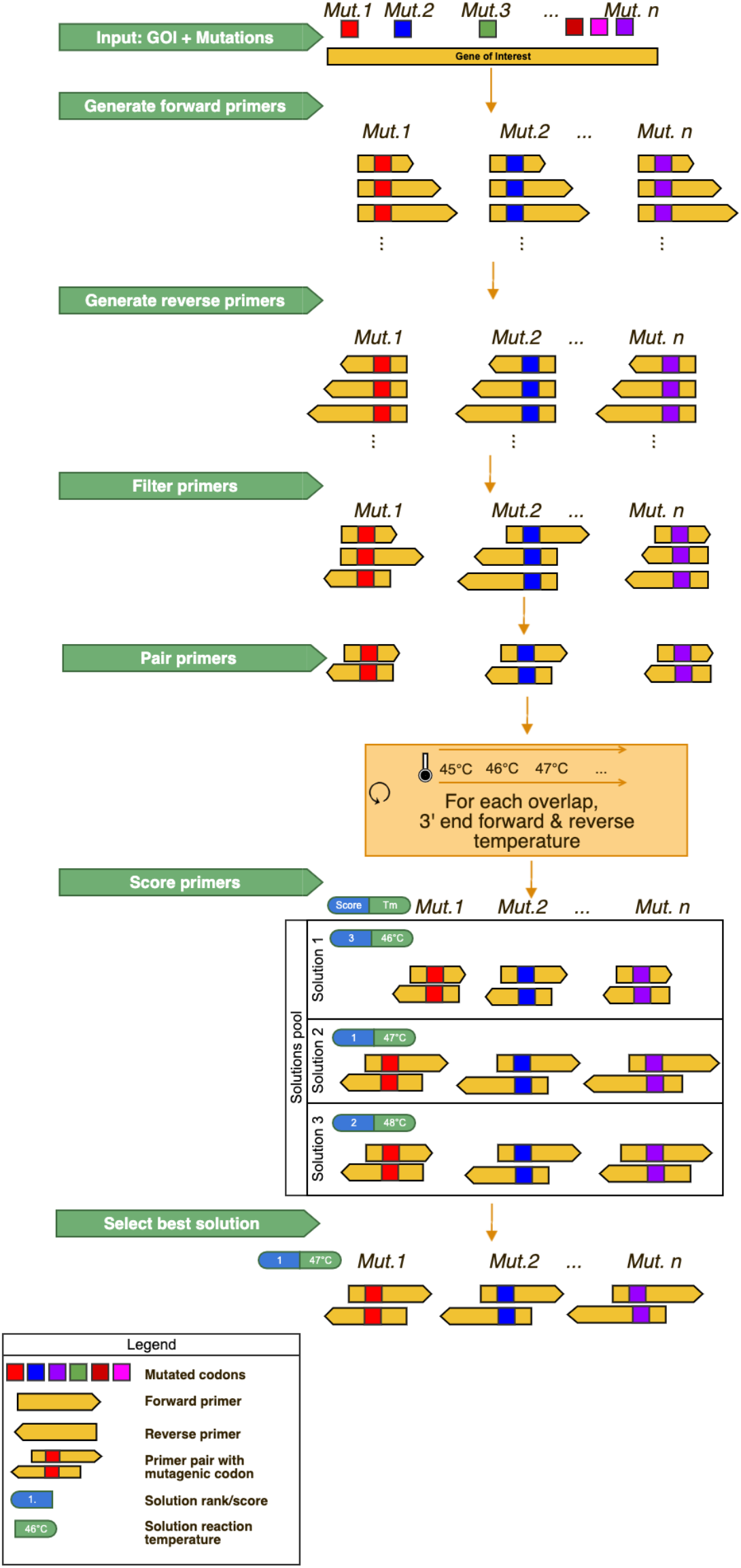
Site-Scanning Saturation Mutagenesis (SSSM) brute-force algorithm for the design of mutagenic primer pairs. In the first step, the algorithm generates all forward and reverse primers for each mutation site via Primer3 binary or independently of Primer3 (depending on user input and selections). The resulting primers are then filtered based on user-specified 3’ end range. Then, the algorithm selects pairs of forward and reverse primers with an overlap size within user-specified range. Given that Primer3 binary automatically filters hairpins, homo and heter-odimers, for each mutation site the algorithm checks if forward and reverse primers could be paired. In case that primer pairs could not be combined for a given site, the algorithm generates primers at all possible offsets around that mutation site in both directions, independently of Primer3 binary and with less stringent conditions (i.e. no hairpins or primer-dimers checks). At the end of the primer generation cycle, the algorithm iterates over a range of reaction temperatures (forward and/or reverse Tm_3’_ and Tm_overlap_ at 3°C and 2°C steps, respectively) and for each pair or triplet of Tm’s and for each mutation site the algorithm selects primer pair with the best cumulative primer pair score. The algorithm scores each primer based on the deviations of primer values from the desired input values of: 3’ length, GC content, overlap and 3’ end Tm’s, and optionally (if specified) also account for the distance of the hairpins and primer-dimers Tm’s from the optimal 3’ end Tm (see Materials and Methods). Finally, the algorithm produces a pool of possible solutions for each reaction temperature from which it selects the best scoring set of primer pairs based on a cumulative solution score. Typically, SSSM primers carry ‘NNK’ codons that encode for all 20 amino acids, degenerate codons can be specified directly or generated on the fly by a heuristic degeneracy algorithm,

**Supplementary Figure S2.**
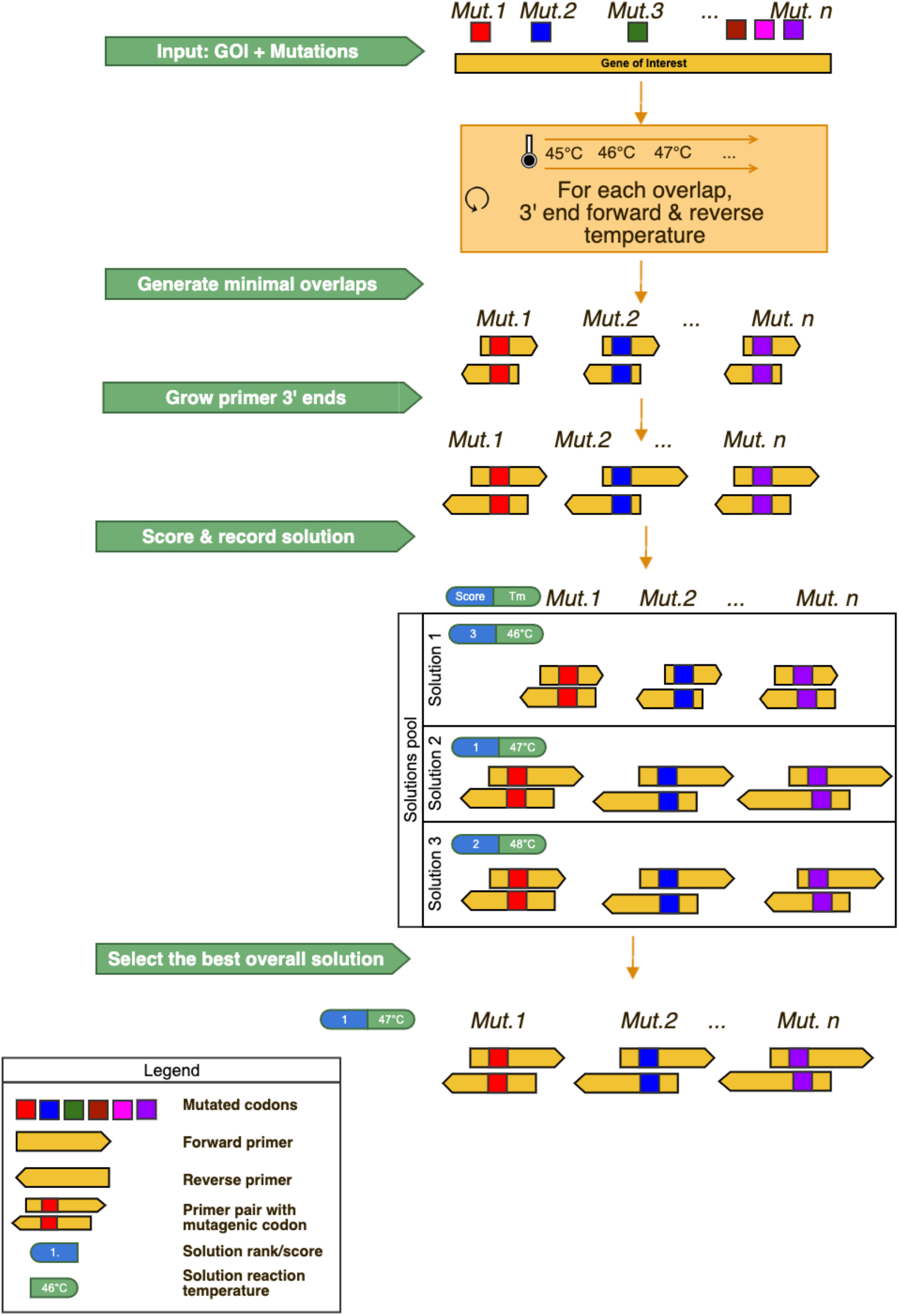
Site-Scanning Saturation Mutagenesis (SSSM) fast-approximation algorithm for the design of mutagenic primer pairs. The algorithm takes into consideration multiple parameters such as primer pair overlap size, melting temperatures of overlap and 3’ end of primer etc. At first, the algorithm iterates (3° C at a time) through a user-specified range of overlap melting temperatures and for each mutation site it finds the shortest primers’ overlap with a melting temperature (Tm_overlap_) that is within half of the allowed temperature deviation between overlaps and 3’ ends (Tm_diff_). Once minimal primer overlaps are identified, the algorithm iterates (2° C at a time) through the user-specified range of the melting temperatures of the 3’ ends (Tm_3’_) of forward and/or reverse primers (independently or combined, as specified) and through the specified range of overlap melting temperatures (3° C at a time, Tm_overlap_). Then, for each mutation site the algorithm selects the primers’ overlap from step 1 that has the closest overlap melting temperature to the current overlap melting temperature (Tm_overlap_). To achieve the desired 3’ end temperature (Tm_3’_), the algorithm iteratively grows the forward and reverse primers (one base at a time) from the overlaps outwards until the desired Tm3’ is reached. Once, primer pairs are grown and reach the desired Tm, the algorithm scores and ranks them based on their deviations from expected values of: 3’ length, GC content, overlap and 3’ end Tm’s, and optionally also account for the distance of the hairpins and primer-dimers Tm’s from the optimal 3’ end Tm (see Materials and Methods). This procedure results in a candidate solution for each iterated reaction temperature, where each solution consists of set of mutagenic primer pairs for all specified mutation sites. At its last step, the algorithm scores all solutions and select the best solution. Typically, SSSM primers carry ‘NNK’ codons that encode for all 20 amino acids, degenerate codons can be specified directly or generated on the fly by a heuristic degeneracy algorithm,

**Supplementary Figure S3.**
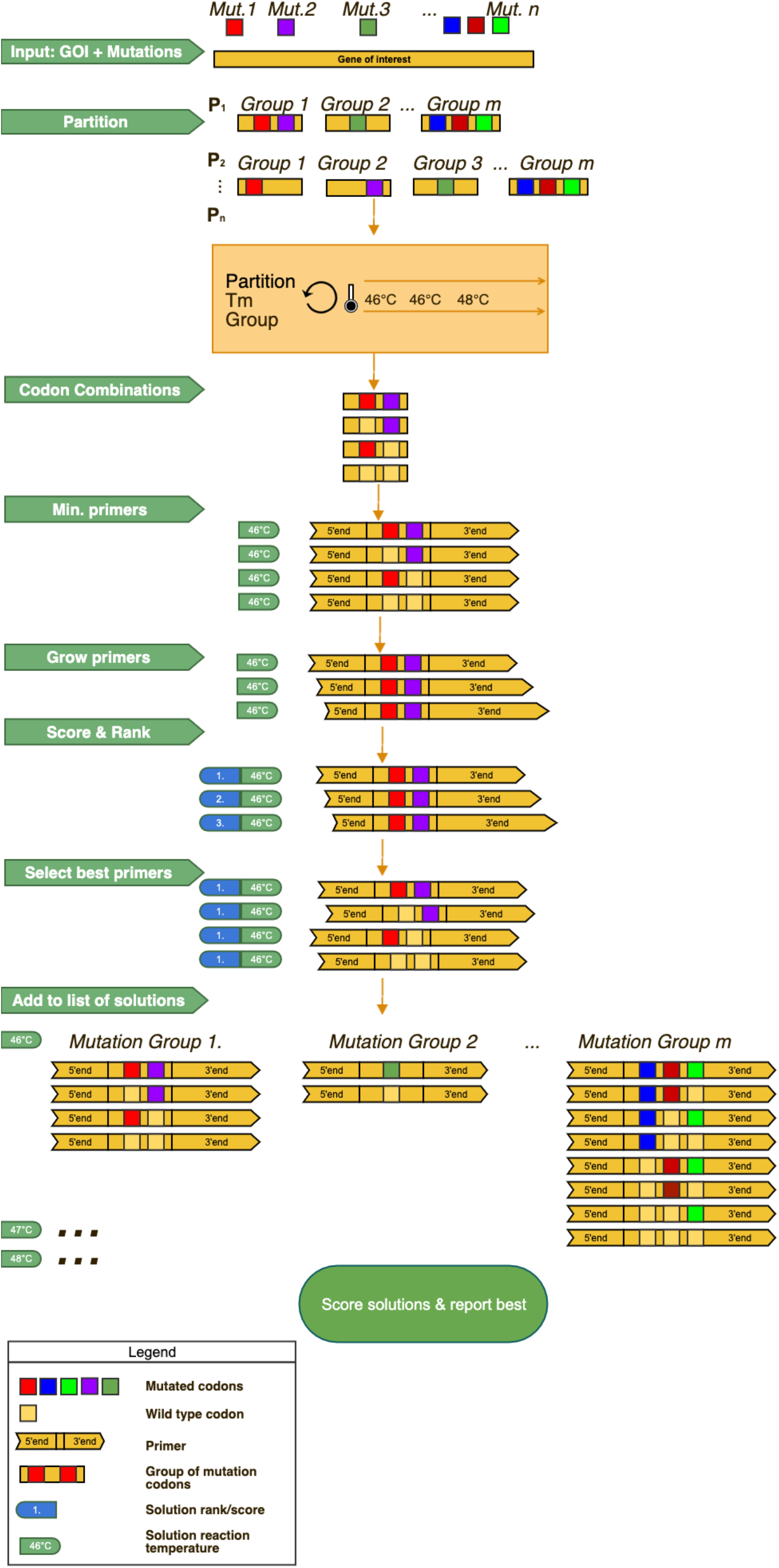
Mutation Maker’s algorithm for Multi Site-Directed Mutagenesis (MSDM) workflow using the QuikChange Lightning Multi Site-Directed Mutagenesis kit (QCLM). To reduce complexity and cost, the MSDM workflow outputs degenerate (default) primers that each may cover one or more mutation sites. Depending on the selected options, primers may be used simultaneously (non-overlapping option) or in successive rounds of MSDM experiments (default). Importantly, in both overlapping and non-overlapping modes Mutation Maker outputs unidirectional primers that never overlap adjacent mutation sites. The algorithm first finds all possible partitions of mutation sites, each consists of groups of mutation sites that can be covered by a single primer, given the input constraints. Then, the algorithm iterates through all partitions (P_1_…P_n_), the entire primer Tm range (1° C at a time) and through all mutation groups. During this cycle the algorithm computes all codon combinations for each mutation group using random codons that encode the desired amino acids (excluding rare codons) and the original parental codons. Thereafter, the algorithm creates minimal primers for each mutation combination while considering all possible offsets around each mutation site. Next, the algorithm extends the 3’ ends of these minimal primers (one base at a time) until they reach the desired Tm at each Tm step. These primers are then scored and ranked, and the best primer for each codon combination in each group is selected. The solution is then added to the list of candidate solutions. Once the algorithm finished looping through all partitions, the entire Tm range and all mutation groups, each solution is scored, and the best solution is ultimately reported. The MSDM cumulative score is synthesized from deviations from the optimal designed criteria values and from the fraction of mutations that are covered by the primer set. If a check for homo- and hetero-dimers is selected by the user, the algorithm scans the list of candidate solutions for primer-dimers among all possible primer pair combinations in each primer set and penalizes the solution score accordingly. To guide even incorporation of mutations, primer premix ratios are also computed for each site, but only for the best reported solution.

**Supplementary Figure S4.**
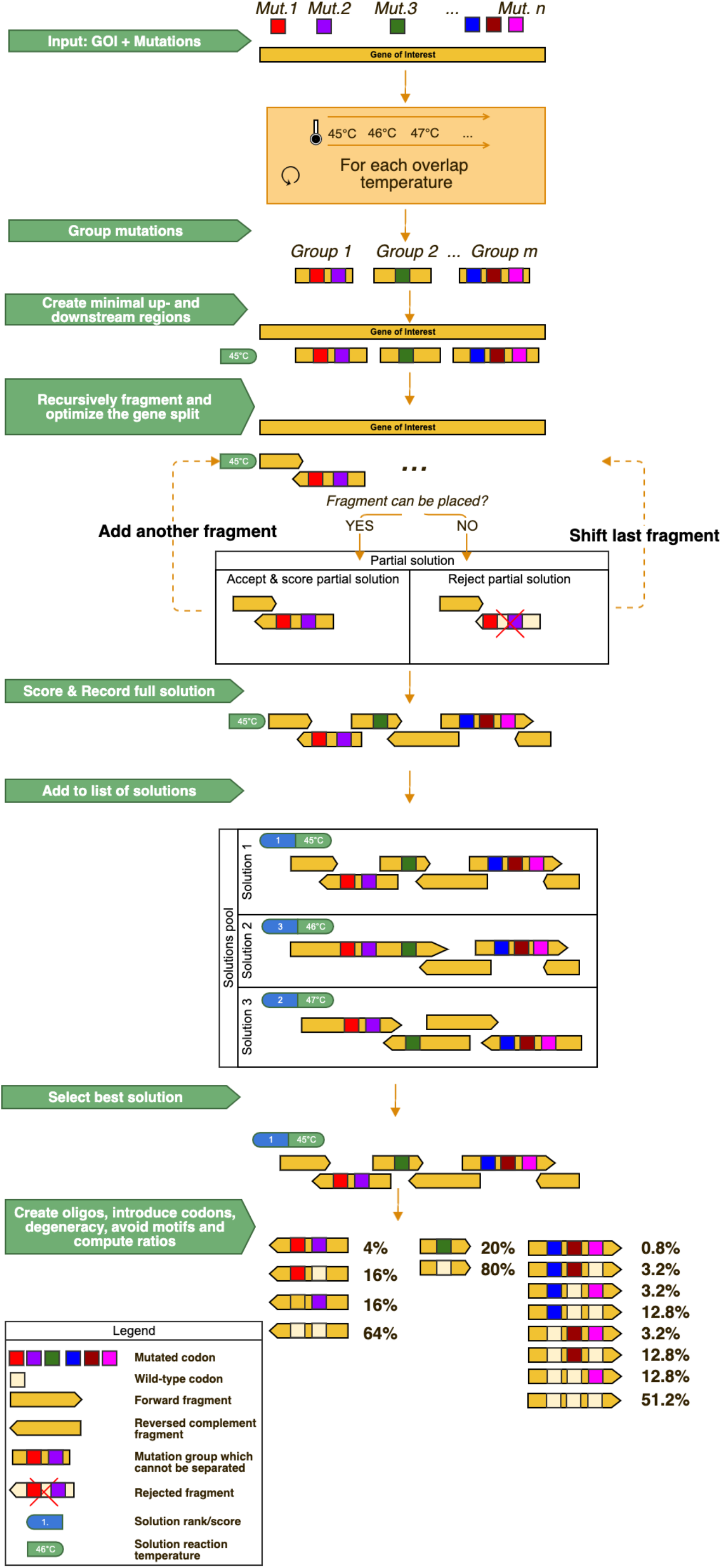
Mutation Maker’s gene synthesis algorithm for mutagenic oligos design using the PCR-based Accurate Synthesis (PAS) protocol. The PAS workflow splits a given gene sequence into an even number of overlapping mutagenic oligos in alternating orientations (forward and reverse complement) that have consistent length and overlap Tm properties. To do so, the algorithm iterates 1°C at the time through the user-specified Tm range for fragment overlaps and first groups the mutations that must reside on the same gene fragment. It does so simply by checking whether the sequence between these mutations can include the overlap between two consecutive PAS fragments given the specified constraints i.e. if the sequence Tm is not lower than the minimal overlap Tm (at that Tm iteration) and if its length is higher than the minimum overlap length. Then, around each mutation group the algorithm generates minimal up- and down-stream sequences that act as guiding constraints for gene fragmentation and can potentially serve as overlaps between two consecutive PAS fragments. To create these potential overlaps, the algorithm extends these sequences (one base at a time) until the resultant extension Tm’s reach to the current iteration overlap Tm and respect the overlap size constraints. At the next step, a backtracking algorithm recursively fragments and optimizes the split of the gene by optimizing the lengths and the positions of PAS fragments or by adding, if necessary, new mutation-less fragments until the entire gene sequence is covered. The backtracking algorithm adds one fragment at the time (starting from the first nucleotide of the gene) and placing it at the first possible position given the problem constraints. When the next fragment cannot be placed due to constraint violation the algorithm shifts the last fragment. If the last fragment cannot be shifted any more (due to constraints violation) the algorithm discards the last fragment and tries to change the previous fragment and so on. In each step, this process creates partial solutions of the gene split that are evaluated using a simplified scoring function that considers only the deviation from the optimal fragment length and hairpin and homodimers Tm’s. Importantly, partial solution scores are recorded only for performance optimization purposes in order to avoid exhaustive search across all possible partial solutions. If the algorithm reaches a feasible solution that covers the entire gene sequence, it computes a complete solution score that considers deviation from all optimal values. However, it records this solution only if its score is better than previously recorded score for the same melting temperature step. The backtracking algorithm continues until it finishes its search or until it reaches a timeout of 5 seconds for each overlap Tm iteration. Once the algorithm completes iterating through the entire user-specified overlap Tm range, the best scoring set of gene fragments is stored. At its final stage, the algorithm introduces the codons to these fragments in order to produce the mutated oligos. If mutations are specified at the amino acid level, the algorithm randomly picks codons (not rare codons) for each mutation site only if they respect the motifs exclusion constraint. If substitutions are specified at the DNA level, the algorithm introduces the exact codon changes requested unless undesired motifs are created. In case that multiple mutations reside on the same fragment, the algorithm outputs all possible oligo variants. The gene synthesis workflow also invokes the degeneracy algorithm when the user specifies multiple substitutions at a single site and opts for degeneracy. At the very end, the algorithm computes the oligo ratios in the reaction mixture that are necessary to achieve the user-defined mutation frequency in the library.

